# Structure-Infused Protein Language Models

**DOI:** 10.1101/2023.12.13.571525

**Authors:** Daniel Peñaherrera, David Ryan Koes

**Affiliations:** University of Pittsburgh

## Abstract

Embeddings from protein language models (PLM’s) capture intricate patterns for protein sequences, enabling more accurate and efficient prediction of protein properties. Incorporating protein structure information as direct input into PLMs results in an improvement on the predictive ability of protein embeddings on downstream tasks. In this work we demonstrate that indirectly infusing structure information into PLMs also leads to performance gains on structure related tasks. The key difference between this framework and others is that at *inference time* the model does not require access to structure to produce its embeddings.

## Introduction

Protein embeddings have much to offer *in silico* methods specialized for predictive tasks that are relevant for drug discovery such as binding site identification, lead optimization, and property prediction. Furthermore, protein embeddings offer a computational convenience as they obviate the need for feature engineering. Multi-modal foundation models for protein biology compress information from both sequence and structure data [1, 2]. Current multi-modal methods, however, require access to both the sequence and structure at inference time. This is a limitation to be avoided, especially for next-generation multi-modal models that aim to incorporate information of protein sequence, structure, and dynamics.

Here we circumvent this requirement by engineering a protein language model that is forced to infuse information about a protein’s 3D structure during *training*, yet only requires sequences as input at *inference* time. We show that embeddings from a PLM trained in this way achieve better performance on structure-related downstream tasks than a sequence-only baseline.

## Methods

Structure-infused protein language models (SI-PLM’s) are composed of non-interacting sequence and structure modules, as shown in Figure 1. The sequence-only module is a standard transformer model [3] whereas the structure module is a Geometric Vector Perceptron graph neural network (GNN) [4] followed by several layers of multi-head attention. We featurize amino acid sequences as tokens and their corresponding 3D structure as a set of geometric scalar and vector quantities. We construct a K-nearest neighbor graph from the protein structure, where nodes are at the *C*_*α*_ positions and have scalar and vector features representing the atoms of the corresponding residue and edge features are based on distances between neighboring nodes.

**Figure 1:**
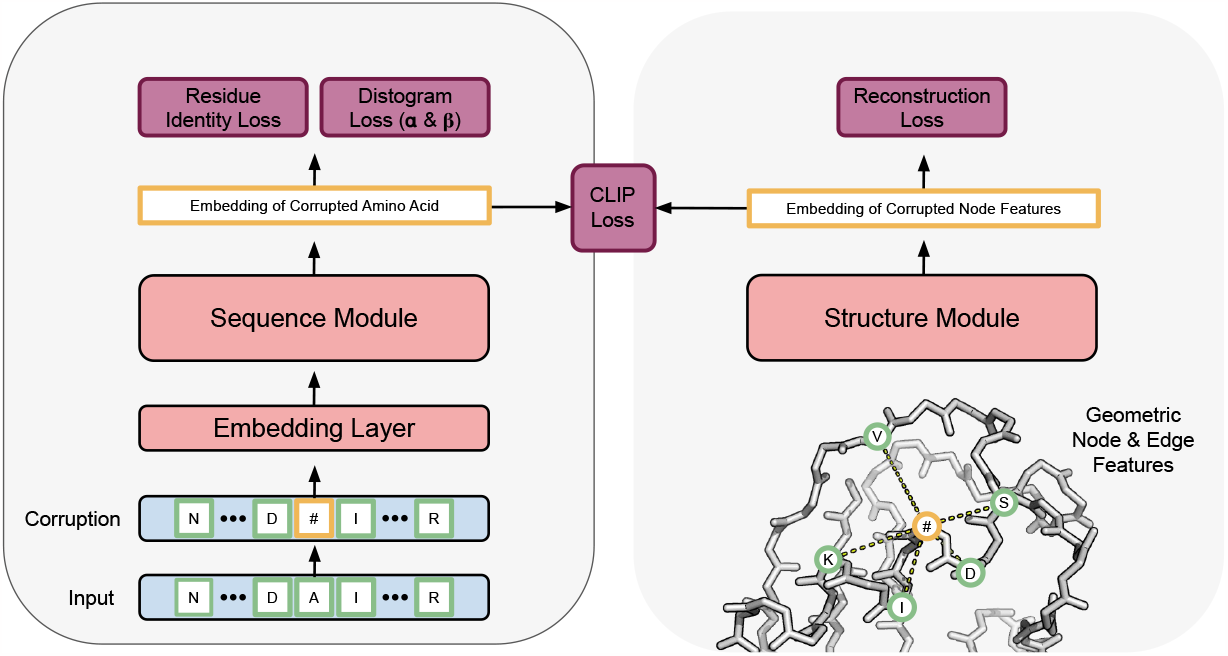
During training the sequence model is encouraged to infuse information about 3D structure into its hidden representations. At inference time, the structure module along with the prediction heads are discarded. One only requires the protein sequence to obtain the embeddings.

During training 15% of the amino acids in a given sequence are chosen to be masked or replaced, as is conventional in BERT-style pre-training [5]. The corresponding node features in the GNN are corrupted to isotropic Gaussian noise. The sequence and structure modules separately process the corrupted input in an attempt to denoise the data. We apply a cross-entropy loss to the sequence module for predicting the corrupted amino acid identity and employ as mean-squared-error loss to the structure module for reconstructing the original node features. Additionally, the embeddings of corrupted amino acids and corrupted node features are used to compute a contrastive loss (CLIP)[6]. Lastly, because the protein structures are available during training, we identify the corrupted amino acids’ nearest neighbors and incorporate a distogram loss into the training of the sequence module.

Massachusetts Institute of Technology - Molecular Machine Learning Conference 2023.

### Dataset

To train our models, we have procured a dataset comprised of X-ray crystallographic structures [7] from the PDB and Alphafold2 predicted structures of the Swiss-Prot sequence database [8], a combined total of 566,118 protein structures. Training and validation splits were generated by clustering sequences at a 90% sequence similarity cutoff using MMSeq2[9]. To evaluate our models, we gathered datasets for three tasks - 3 class secondary structure prediction [10], binary classification of binding residues [11], and 10 class classification of protein localization (DeepLoc) [12].

### Evaluation

To evaluate SI-PLM, we first trained a baseline sequence-only PLM on the same sequences from our dataset. Weights from the sequence encoders of both the SI-PLM and baseline PLM were frozen and used to generate amino acid embeddings. These were provided as input to a small 2-layer convolutional neural network to predict classes for either the secondary structure task or the binding residue task. In the case of DeepLoc, a light attention network, as detailed in [12], was used. We additionally trained these models on a one-hot encoding of the sequences for comparison.

## Results

Table 1 lists the average per-class accuracy for two of the three benchmark tasks, secondary structure and DeepLoc. For the binary prediction task of predicting whether a given residue binds to either a small molecule, metal ion, or nucleic acid, the majority of residue do not bind. This results in a large class imbalance and so we opt to report the more informative F1-score. Embeddings from SI-PLM exhibit the best performance for all but one test set and the largest improvement in performance is seen for the most structure-related task (secondary structure prediction). In all cases the models trained using embeddings outperform a simple one-hot representation. Overall these results indicate that the proposed training strategy for a SI-PLM does indeed improve performance over the sequence-only baseline and one-hot based model.

**Table 1:**
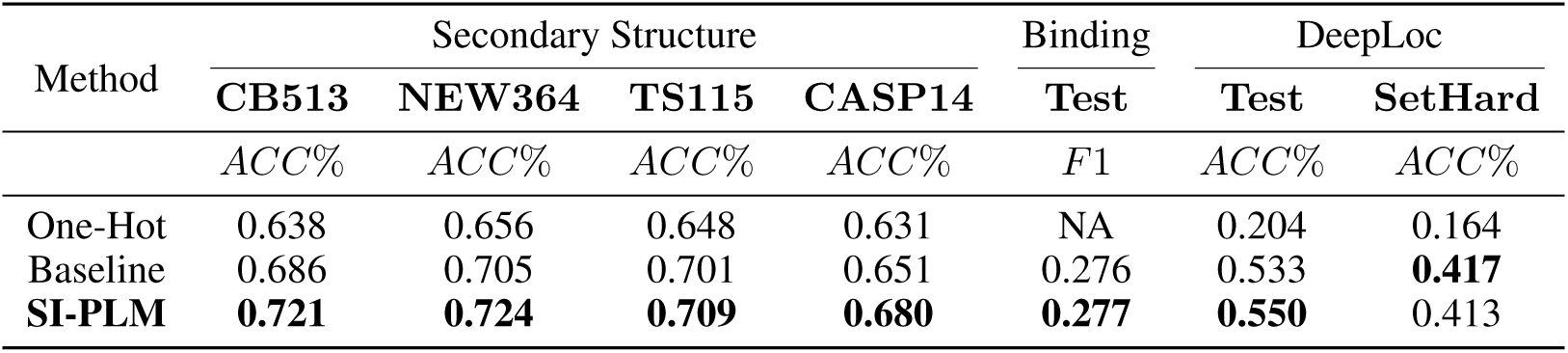
Transfer Learning Performance Across Three Benchmark Tasks.

| Method | Secondary Structure |  |  |  | Binding | DeepLoc |  |
| --- | --- | --- | --- | --- | --- | --- | --- |
|  | CB513 | NEW364 | TS115 | CASP14 | Test | Test | SetHard |
|  | *ACC%* | *ACC%* | *ACC%* | *ACC%* | *F1* | *ACC%* | *ACC%* |
| One-Hot | 0.638 | 0.656 | 0.648 | 0.631 | NA | 0.204 | 0.164 |
| Baseline | 0.686 | 0.705 | 0.701 | 0.651 | 0.276 | 0.533 | **0.417** |
| **SI-PLM** | **0.721** | **0.724** | **0.709** | **0.680** | **0.277** | **0.550** | 0.413 |

## A Appendix

### A.1 Contrastive Loss

We employ the CLIP [6] loss function to enforce that the embeddings coming from the sequence and structure modules are similar to each other by the following:

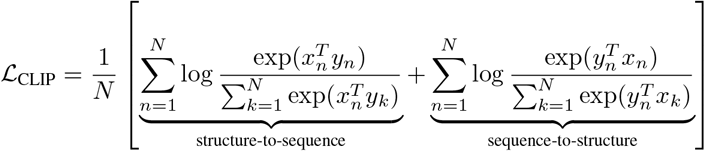

where (*x*_*i*_, *y*_*i*_) ∈ ℝ^*d*^ represent amino acid and node embedding pairs, respectively, and *N* is the number of corrupted amino acids in the batch.

### A.2 Distogram Loss

The sequence module of the SI-PLM leverages a distogram prediction head to predict distance between neighboring residues. This is accomplished by an attention mechanism that computes pair representation *Z*_*ij*_ between amino acid embedding *x*_*i*_ and neighboring amino acids *x*_*j*_∀*j* ∈ 𝒩 (*i*):

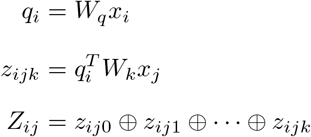

where *W*_*q*_ and *W*_*k*_ are weight matrices and ⊕ denotes the concatenation operator. The pair representations are then projected into 64 distance bins that cover a range from 2 *Å* to 22 *Å*, as is done in AlphaFold2[13]. The actual loss is calculated as the cross-entropy over all neighbors of a selected amino acid:

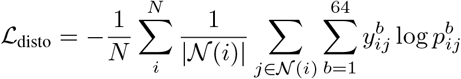

where *N* is the number of corrupted amino acids in a batch, | 𝒩 (*i*)| is the number of neighbors for a given node *i*, 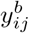 are the target binned distances, and 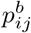

